# Ang2 and TAT targeting of leptomeningeal disease by the intravenous and intrathecal routes: a comparative analysis

**DOI:** 10.64898/2026.06.29.735336

**Authors:** Chung-Fan Kuo, Tobi Babayemi, Kha Uyen Dam, Shaokuan Zheng, Hongwei Yang, Rachael W. Sirianni

## Abstract

Leptomeningeal disease (LD) refers to the metastasis of cells to the leptomeningeal membranes in the central nervous system (CNS), and it is a deadly complication of several different types of cancer, including breast cancer (BC) and pediatric medulloblastoma (MB). Angiopep-2 (Ang2) and transactivating transcriptional activator (TAT) peptides have been reported to transport therapeutic cargos into the CNS. Here, compared the LD-targeting capability of Ang2 vs TAT by intrathecal (intracisternal magna, ICM) vs intravenous (IV) routes of administration. We first generated two xenograft models of LD by directly infusing breast cancer cells (MDA-MB231) or medulloblastoma cells (HDMB03) via the ICM route, characterizing models for survival, tumor growth patterns, and presence of hydrocephalus. Second, we administered fluorescently labeled Ang2 or TAT peptides either IV or ICM into tumor bearing mice to examine ligand localization. We discovered that the median survival of both models was negatively related to the number of the cells infused. While HDMB03 cells tended to metastasize preferentially to the brain region, MDA-MB231 cells tended to metastasize preferentially to the spinal cord. Compared to the healthy control, MB-LD yielded a 7.3-fold increase and BC-LD a 26.5-fold increase in ventricular volume. Furthermore, targeting achieved by TAT was significantly higher than targeting achieved by Ang2 in thoracic spine for the MB-LD model. For BC-LD, TAT targeting was superiot to Ang2 in the olfactory bulbs, brain stem, thoracic spine, and lumbar spine regions. Significantly, these data provide evidence that ICM will be a preferable route of administration over IV for maximally targeting LD.

## 1. Introduction

Leptomeningeal disease (LD), also known as leptomeningeal metastasis, refers to the dissemination of a primary tumor across the leptomeninges that cover the brain and spinal cord, and it is diagnosed when tumor cells are detected in cerebrospinal fluid (CSF) [1, 2]. Several primary tumors have been reported to highly associated to LD development, such as melanoma, lung cancer, pediatric brain cancer, and breast cancer [3]. A diagnosis of LD carries very poor prognosis and is considered fatal in a recurrent setting. One of the major challenges for treatment of LD is that it cannot be readily surgically resected, and disseminated disease is often poorly responsive to traditional chemotherapy.

Targeted therapy is a promising new approach to improve overall survival for LD patients [4]. While molecular targeting is focused on tackling the specific molecular mutations in specific type of cancer [4], peptide-ligand targeting is addressed to tackle the tumor cell surface molecules in order to selectively destroy malignant cells without harming healthy cells [5]. Targeting offers several opportunities to innovate on new therapeutic approaches in LD, enabling potentially selective delivery of therapeutic agents (small molecules, antibodies, oligonucleotides, and drug loaded nanoparticles) to malignant tissues while decreasing toxicity to off-target tissues. Unsurprisingly, the use of peptide, nanobody, and antibody ligands are a major focus for enabling transport of drugs or drug carriers across the blood-brain barrier (BBB) to reach the central nervous system (CNS) [6, 7].

Here, we focus on Angiopep-2 (Ang2), which is a 19-amino-acid peptide that possesses well-validated ability to achieve transcytosis across the endothelial cells that comprise the BBB through its selective binding with LRP-1 receptors; Ang2 has found success in both preclinical and clinical settings [6, 8]. As an alternative, we also examined the transactivating transcriptional activator (TAT), which is a widely used cell-penetrating peptide [9]. TAT rapidly enters cell through electrostatics interaction between TAT’s positively-charged amino acids (arginine and lysine) and negatively-charged phospholipids on the cellular membrane [10], and this capability has also been useful for targeting the BBB. Both Ang2 and TAT have been widely applied to brain delivery. For example, intravenously (IV) administered therapeutics linked to Ang2 demonstrated efficacy against primary brain tumors [11, 12], brain metastatic lung cancer [13], and brain metastatic breast cancer [14] in pre-clinical studies. Several phase I and II clinical trials were also designed to treat different brain cancers using a paclitaxel-Ang2 conjugate (ANG1005) (NCT00539344, NCT00539383, NCT02048059, and NCT01967810). Similarly, TAT was able to deliver therapeutic molecules and drug-carried nanoparticles to brain metastases in breast cancer [15, 16].

Although some success has been had for use of Ang2 for treatment of LD by the IV route, here, we were interested to explore an alternative route of administration. Intrathecal (IT) drug delivery involves the infusion of therapeutic agents directly into the CSF via lumbar, ventricular, or cisternal access points. IT administration enables substances to directly access the subarachnoid space (SAS) of the CNS, where they mix with CSF and flow across LD that is exposed on the surfaces of the brain and spinal cord [17]. We and others have shown that IT administration results in very high levels of drug within the CSF while minimizing systemic exposure [17, 18].

Despite the high level of activity in developing therapeutics for both potential routes of administration (IV, IT), relatively few studies have examined the comparative benefits of targeting by one vs the other strategy. With this long-term goal in mind, we sought to generate models of LD that would enable us to systematically examine feasibility of targeting malignant cells that have metastasized to the leptomeninges with Ang2 or TAT. This work focuses on two cancer models (triple-negative breast cancer (TNBC) and Group 3 medulloblastoma (MB)) as the primary tumor source due to the clinical significance of the disease [19, 20]. We developed these models and administered fluorescently labeled Ang2 and TAT peptides by the IV and IT routes, enabling us to rigorously examine the potential advantages of different drug delivery approaches for treatment of LD.

## 2. Materials and methods

### 2.1. Material

Avantor® Seradigm, Select Grade Fetal Bovine Serum (FBS) was purchased from Avantor (Radnor Township, PA). HyClone Penicillin-Streptomycin 100X solution was obtain from Cytiva (Marlborough, MA). MEM Nonessential Amino Acids Solution (100X) was bought from Corning (Corning, NY). MDA-MB-231/GFP-luciferase stable cell line was purchased from Fenicsbio (Halethorpe, MD). Ethylenediaminetetraacetic Acid, Disodium Salt Dihydrate was received from Fisher Scientific (Waltham, MA). TAT-FITC, TAT-Cy5, TAT-Cy3, Ang2-FITC, Ang2-Cy5, and Ang2-Cy3 was custom synthesized by GenScript (Piscataway NJ). DPBS (no calcium, no magnesium), Trypsin (2.5%, no phenol red), Dulbecco’s Modified Eagle Medium (DMEM, high glucose, pyruvate), and RPMI 1640 Medium were received from Gibco (Grand Island, NY). ProLong™ Glass Antifade Mountant and DAPI and Hoechst Nucleic Acid Stains were obtained from Invitrogen (Waltham, MA). NOD.Cg-Prkdcscid Il2rgtm1Wjl/SzJ (NSG) was purchased from Jackson Laboratory (Bar Harbor, ME). Sucrose, lyophilization certified was received from Ops Diagnostics (Lebanon, NJ). IVISbrite D-Luciferin Potassium Salt Bioluminescent Substrate was bought from Revvity (Waltham, MA). Pierce™ Firefly Luciferase Glow Assay Kit and Paraformaldehyde (16% w/v aq. soln., methanol free) were received from Thermo Scientific (Waltham, MA)

### 2.2. Cell culture

GFP and luciferase transduced, MDA-MB231-GFP/Luc, representing human TNBC, were purchased from Fenicsbio Halethorpe, MD. MCherry and luciferase transduced, HDMB03- mCherry/Luc, representing human MB, were generously provided by Dr. Robert Wechsler-Reya (Columbia University, NY, USA). MDA-MB231-GFP/Luc cells were maintained in DMEM with 10 % FBS and 1% penicillin-streptomycin. HDMB03-mCherry/Luc were maintained in RPMI1640 with 10 % FBS, 1% penicillin-streptomycin, and 1% nonessential amino acids. All cells were incubated at 37 °C with 5 % CO_2_.

### 2.3. *In vitro* luciferase assay

MDA-MB231-GFP/Luc cells and HDMB03-mCherry/Luc cells with a desired cell number were collected after trypsinization. Cells were centrifuged at 1400 rpm for 4 mins, and the supernatant was carefully removed. Fifty microliters of 1x D-Luciferase assay buffer per condition was added to the cell pellets. The cell-contained luciferase assay buffer was then transferred into a 96-well plate, followed by the luminescence intensity measurement using a microplate reader (Tecan Spark, Männedorf, Switzerland) at 20 minutes incubation time.

### 2.4. Tumor induction, monitoring, and tissue processing

All experiments, procedures and animal care practices were approved by the UMass Chan Medical School’s Institutional Animal Care and Use Committee and were performed in accordance with all relevant guidelines. Healthy, female NSG albino mice with age of within 6 to 8 weeks old were used for *in vivo* experiments. The injection procedure was modified from a report published previously [21]. Briefly, on the day of the procedure, animals were weighted prior to the injection. Mice were anesthetized with 2% isoflurane and positioned with the superior aspect of the neck hyperextended to expose the back of the neck. The fur at the top of the head and back of the neck was shaved to expose skin above the cisterna magna. Percutaneous injection was accomplished between the base of the skull and C1 vertebrae with a 10 µL Hamilton neuro-syringe connected to a 33-gauge needle with stopper positioned at 3.5mm (**Figure 1**). Cells were slowly infused into the cisterna magna over 60 to 70 seconds, keeping the needle in place for an additional 5 to 10 seconds before it was withdrawn.

**Figure 1.**
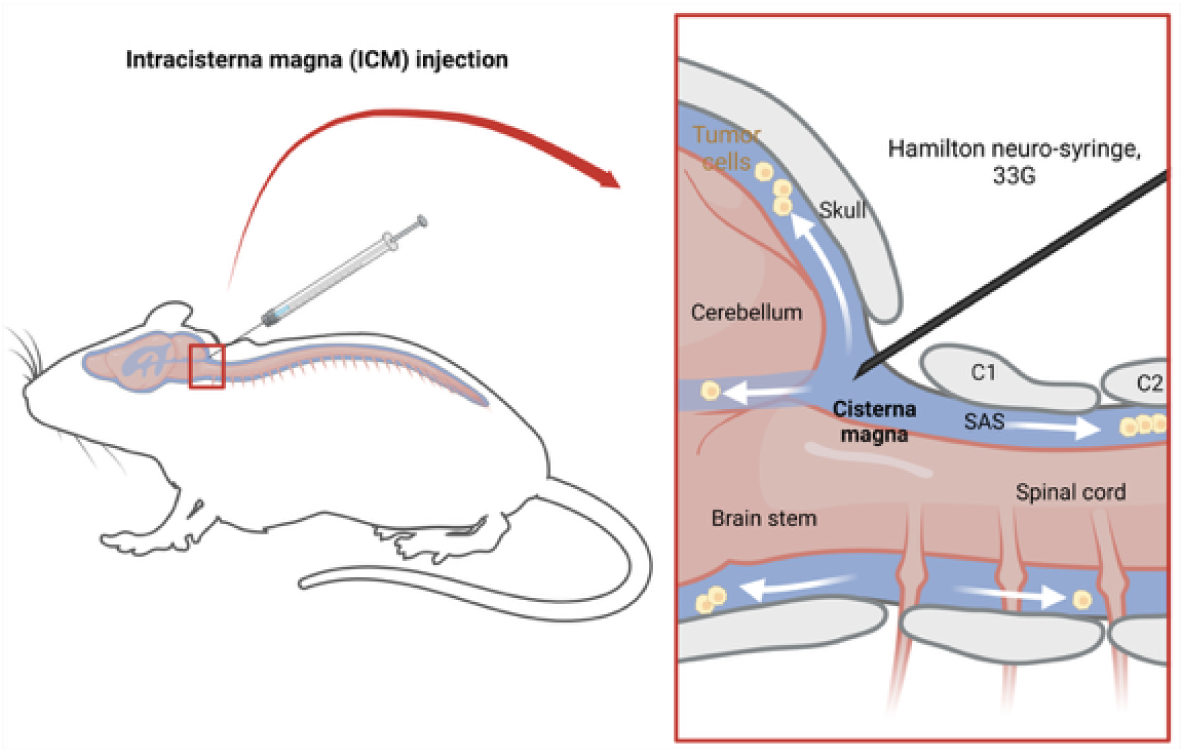
Schematic of the intracisternal magna injection. Tumor cells were injected into mouse’s cisterna magna using a Hamilton neuro-syringe. Injected cells would then have the access of whole SAS to mimic the LD.

Following cell transplant, mice were monitored daily for gross behavior and weight. The growth of the tumor was assessed by In Vivo Imaging System (IVIS) twice per week. Briefly, D-luciferin solution at a concentration of 150 mg/kg was injected into the mice via the intraperitoneal (IP) route. Luminescence intensity was captured by IVIS Spectrum CT (Perkin-Elmer, Waltham, MA) 20 minutes after administration of luciferin substrate. Two regions of interest (ROI) were drawn to collect total flux of the brain and spinal cord. The size of the ROIs was kept identical for all images analyzed. For peptide localization studies, mice were euthanized immediately after IVIS imaging. For the survival studies, mice were euthanized when the mice met the euthanasia criteria (weight loss of >20% and/or presence of any neurological symptoms). Mice were euthanized by transcardial perfusion with heparinized PBS followed by 4% paraformaldehyde. The mouse’s skin, organs, and muscles were carefully removed to isolate the brain and spinal cord-containing skull and vertebral column. These intact tissues were decalcified in 10% EDTA at 4 °C for 14 days and dehydration in 30% (w/v) sucrose solution. Brain and spinal cord tissues were then cryo-protected in OCT and were slied to a thickness of 12 µm.

For tissue fluorescent images and H&E images, tissue slices were stained either with DAPI or hematoxylin and eosin (H&E). Tissues stained with H&E were imaged using a TissueGnostics TissueFAXS SL Q slide scanning microscope equipped with Pl-D674CU-CYL-07451 Color Camera and 20x 0.75 NA Zeiss Plan Apo objective. Images were taken as extended focus projections in widefield mode with a 3-step z-stack consisting of 0.6 µm steps. Autofocus points were automatically determined across the selected region. Tissues stained with DAPI were imaged using a Leica stereoscope (Leica M205 FA, Wetzlar, Germany).

### 2.5. MRI imaging

Magnetic resonance imaging (MRI) was performed on moribund subjects. T2-weighted MRI images of the brains of the tumor bearing mice were obtained using a Bruker BioSpec 70/30 7T MRI in The Advanced MRI Center at UMass Chan Medical School. Anesthesia was induced by 5% isoflurane and maintained at 1-2% isoflurane for the duration of the experiment, with an oxygen flow rate of 2 L/min. Mice (n=1 each for healthy control, MB-LD, and BC-LD) were imaged with a T2-weighted scan of the head in a BioSpec 70/30 USR horizontal bore MR system (Bruker Corporation, Billerica, MA, USA). Images were cropped to isolate the brain, and the ventricles were identified on the basis of brightness through a custom MATLAB software (R2024b) to create a 3D rendering of the ventricles and estimate volume in µL.

### 2.6. Peptide administration

Peptides were administered to mice bearing a moderate tumor load, i.e., when the brain total flux measured by IVIS imaging reached between 10^9^ to 10^10^ photons/sec; this level corresponds to a moderately high tumor burden. Ang2-FITC and TAT-FITC were prepared at a concentration of 15 nmole in 10 µL in 1x PBS and injected into HDMB03-tumor bearing mice via the IT-CM route. Ang2-Cy5 and TAT-Cy5 were prepared at a concentration of 1 nmole in 10 µL in 1x PBS and injected into MDA-MB231-tumor bearing mice via the IT-CM route. For IV experiments, TAT- FITC was prepared at a concentration of 50 nmole in 100 µL (HDMB03-tumor bearing mice) and TAT-Cy5 was prepared at a concentration of 3.33 nmole in 100 µL (MDA-MB231-tumor bearing mice) via the lateral tail vein. The concentration for IV injection were selected to be below the threshold for toxicity reported by others [22]. Mice were euthanized by perfusion and exsanguination either 2 or 24 hours after peptide injection, after which tissues were removed and processed as described above.

### 2.7. Tissue Imaging

The whole brain and spinal cord were sliced to 10-15um thickness on Superfrost Plus slides, and fluorescent images were taken using a Leica stereoscope (Leica M205 FA, Wetzlar, Germany). The ROI at specific locations of the CNS was drawn to quantify the peptide localization with target tissues. Skull and vertebral columns from healthy mice were also collected to set up the background natural fluorescent signal. Decalcification, dehydration, OCT imbedding, and tissue slicing process were further followed. Tissues were imaged using a Leica confocal microscope (Leica TCS SP5 II, Wetzlar, Germany). Imaging settings were adjusted to control for tissue autofluorescence and maintained at the same level for all experiments. Linear adjustments (gain, brightness, or contrast) were applied equivalently to all images.

### 2.8. *In vitro* cellular uptake by fluorescent imaging

MDA-MB231 and HDMB03 cells were seeded onto poly-L-lysine coated glass coverslips overnight at room temperature, with a seeding density of 0.025x10^6^ cells per coverslip, which were then placed in 6-well plates and covered with 200uL of DMEM and RPMI-1640 for MDA- MB231 and HDMB03 cells, respectively. Following 48 hours of incubation at 37 °C in 5% CO_2_, 10 µM of Ang2-Cy3 and TAT-Cy3 were added to MDA-MB231 cultures and 10 uM of Ang2-FITC and TAT-FITC were added to HDMB03 cultures, followed by 2-hr incubation at 37°C. After incubation, cell media was removed and the cells were washed, fixed with 4% paraformaldehyde for 15 minutes at room temperature, and washed again to remove residual fixative. Cells were stained with DAPI for 10 minutes at room temperature followed by washing. The cell-contained glass coverslips were mounted onto glass slides using a ProLong™ Glass Antifade Mountant. All uptake images were taken by Leica inverted microscope (Leica DMIL LED, Wetzlar, Germany). The fluorescence intensity settings of Cy3, GFP, FITC, and DAPI across two peptide conditions were adjusted to control for tissue autofluorescence and maintained at the same level for all experiments to enable direct, unbiased comparison.

### 2.9. Quantitative analysis of cellular uptake by FACS

Confluent MDA-MB231 and HDMB03 cells were seeded into 12-well plate at a seeding density at 250,000 cells/well. After culturing for 48 hours, media was removed, and peptides were added at a concentration of 1 µM or 10 µM for 0.25, 0.5, 1 and 2 hours at a 37°C, 5% CO_2_ incubator. At the end of the experiment time, cells were then washed twice with 1X DPBS, followed by trypsinization. Cells were collected in 0.5 mL 1X DPBS and analyzed with a BD Symphony A5 flow cytometer. Cellular uptake was quantified by dividing the mean fluorescence intensity (MFI) detected in the peptide-incubated cells by MFI measured for cells that were not incubated with peptide.

### 2.10. Statistical analysis

All data are presented as mean ± SD unless otherwise stated. Statistical analysis was performed using Prism software (San Diego, CA) at an alpha level of 0.05. All graphics were generated on BioRender.com.

## 3. Results

### 3.1. Model Development

To examine peptide targeting capability, we first developed and characterized two models of LD, one representing human, pediatric MB (HDMB03), and the other representing human, adult BC (MDA-MD231) (**Figure 2**). While healthy mice gained weight throughout the experiment period, mice bearing HDMB03 tumors began to lose weight immediately after tumor induction; mice bearing MDA-MD231 tumors maintained their weight for at least 3 weeks followed by rapid weight loss (**Figure 2A**). Median survival was negatively related to the number of the cells infused in both models, where higher cell number yielded shorter medial survival. In detail, the median survival for BC-LD models with 0.5m, 0.05m, 0.01m, and 0.005m cell injections was 29, 34, 41.5, and 48.5 days, respectively. The median survival for MD-LD models with 0.5m, 0.05m, 0.01m, and 0.005m cell injection was 16, 17, 24, and 24 days, respectively (**Figure 2B**). The BC-LD model was thus characterized by longer medial survival compared to MB-LD model.

**Figure 2.**
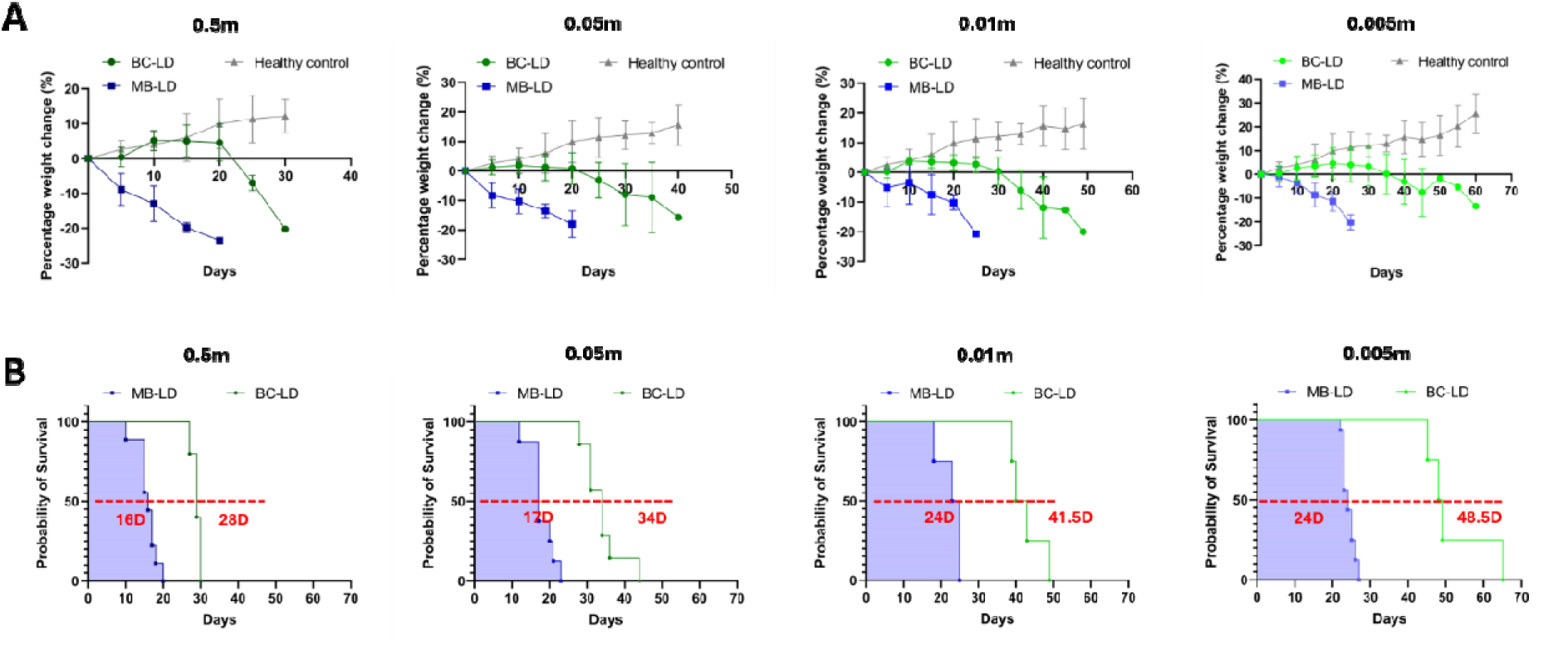
BC-LD model has slower weight drop and longer survival compared to MB-LD model. Weight percentage change (A) and Kaplan-meier survival curve (B) of BC-LD and MB- LD models with different umbers of cells infused, where 0.5m, 0.05m, 0.01m, 0.005m represents 0.5, 0.05, 0.01, 0.005 million cells.

We examined tumor growth in greater detail with IVIS. Cell number was directly and linearly related to measured luminescence *in vitro* (**Figure 3A**). Both LD models developed metastasis across the neuroaxis, reaching both the brain and spinal cord with variable frequency (**Figure 3B-C**). For both models, tumor cells tended to move from the cerebellum region (adjacent to the injection site) to the olfactory bulbs and spinal cord regions, with metastasis frequently observed in the lumbar and sacral regions of the spinal cord. The MB-LD model has relatively higher brain/spinal cord ratios across all cell number groups compared to the BC-LD model (**Figure 3D**). Thus, HDMB03 cells tended to metastasize preferentially to the brain ROI, while MDA- MB231 cells tended to metastasize preferentially to the spinal cord ROI.

**Figure 3.**
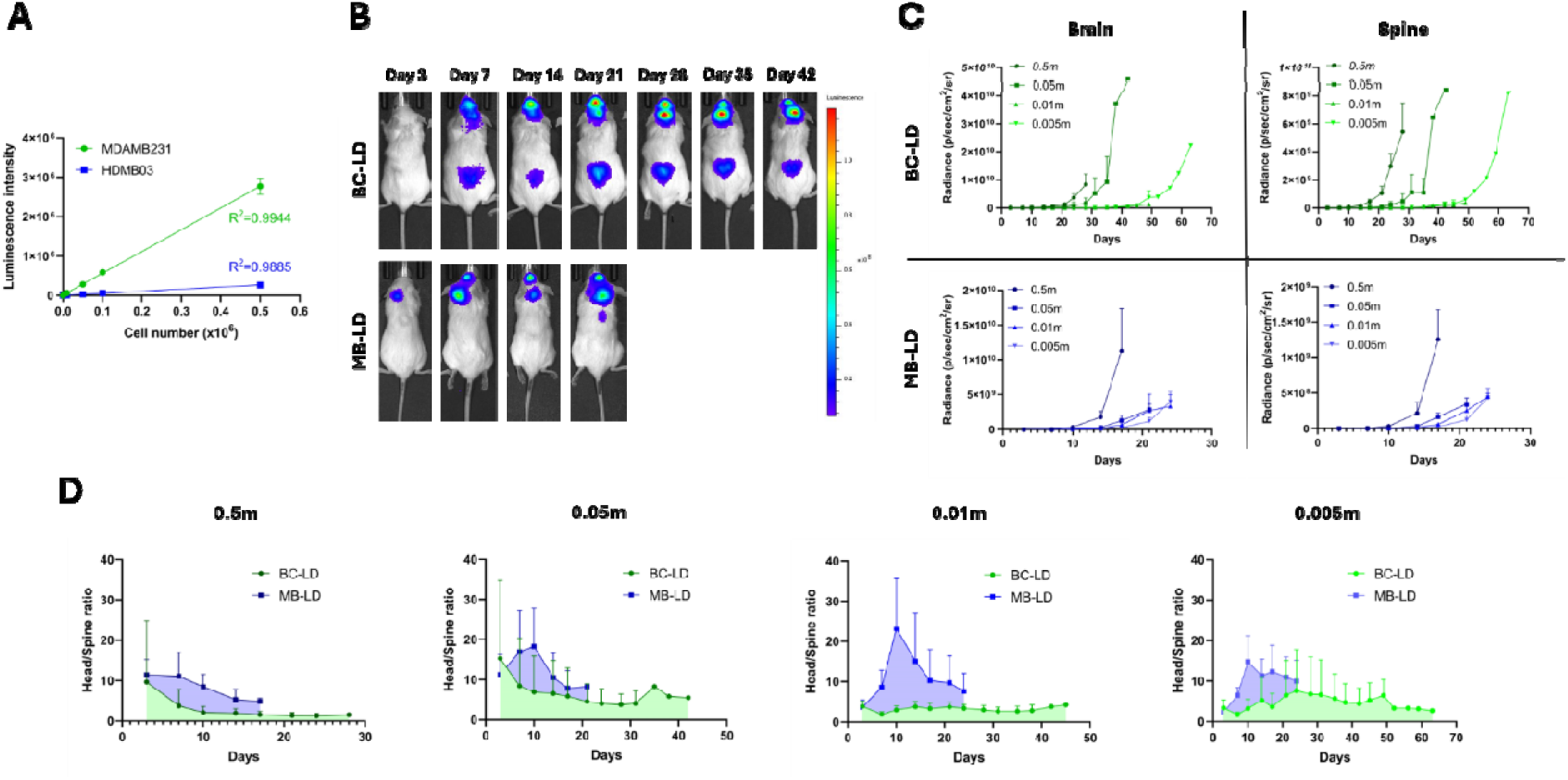
Tumor growth pattern of two LD models was monitored by IVIS system *in vivo*. (A) Linear correlation of the cell number and luminescence intensity of two tumor cell types, measured *in vitro* using a plate reader. (B) IVIS images of two models, selected from one of the mice from 0.005m to represent the group. Animals were imaged twice a week until the day of euthanasia. (C) Tumor growth curves of two models. Two ROIs were drawn to extract the radiance of the luminescence intensity from brain and spinal regions. (D) The luminescence intensity ratio of brain and spinal cord regions of the two models with different cell numbers injected.

We next examined metastasis at the cellular level. For H&E staining, sites of metastasis could be identified by their dark coloration (higher density of cells) and abnormal cellular shape. This is because tumor cells are characterized by a large nucleus, having an irregular size and shape, the nucleoli are prominent, the cytoplasm is scarce and intensely colored [23]. Both LD models exhibited extensive metastasis within the SAS and across the leptomeninges, spanning locations from the olfactory bulbs to the hypothalamus, brain stem, and lumbar spine (**Figure 4A-B**). MDA-MB231 cells are seen to infiltrate into brain parenchyma extensively; however, HDMB03 cells are primarily found in SAS with less evidence of widespread parenchymal infiltration. Both models have strong tumor signals at the ventral side of the brain, especially at the location of hypothalamus. This might be explained by the route of the clearance of the CSF, where CSF flows through the olfactory bulbs and bottom side of the brain then converges into the deep cervical lymph nodes. Ultimately, for both models, the majority of metastatic cells are directly exposed to CSF that flows through the SAS.

**Figure 4.**
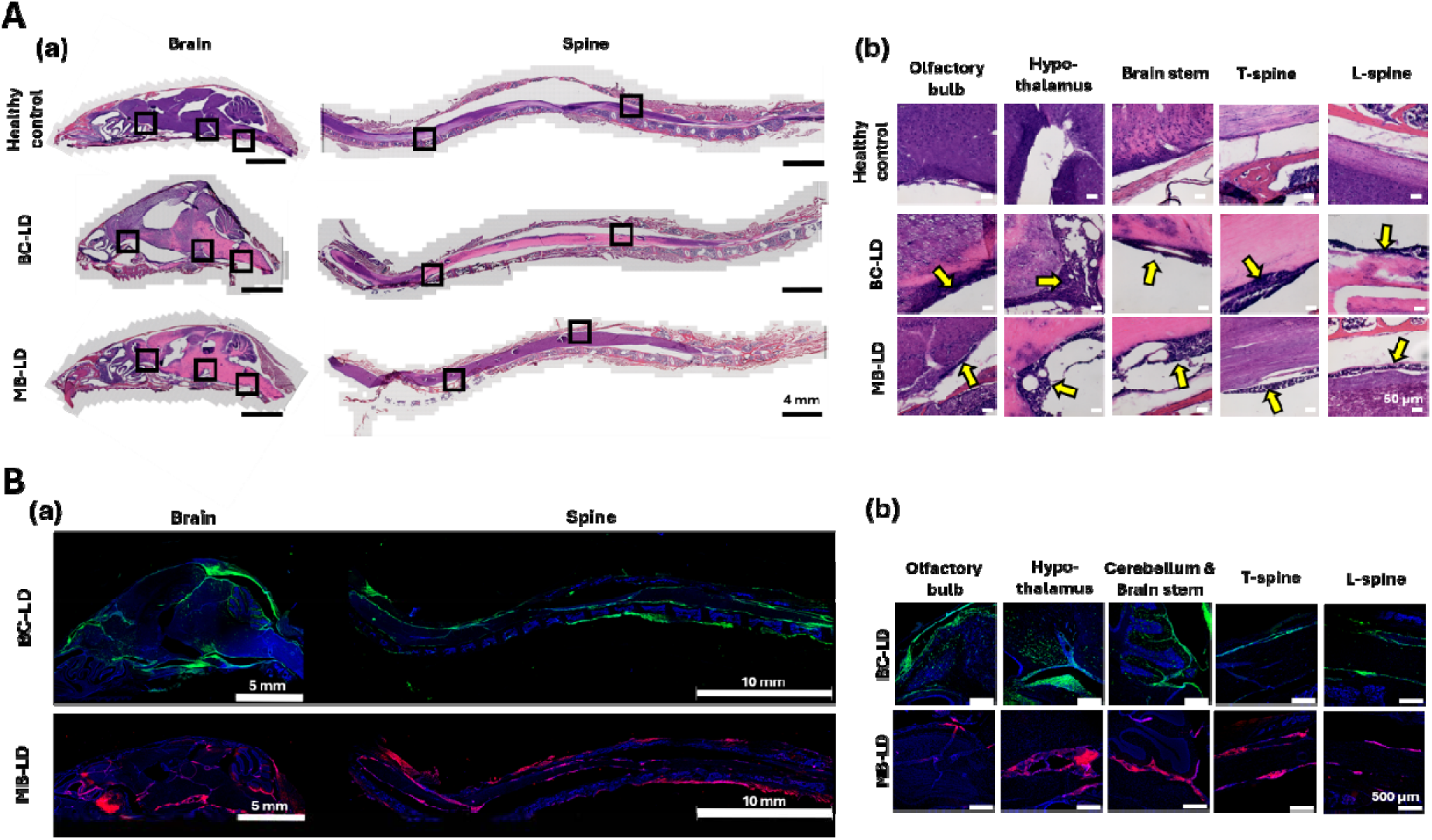
The distribution of the tumor population within the mouse SAS was indicated by H&E staining technique and fluorescence imaging technique. (A, a) The H&E stain images of whole brains and spinal cords of the two LD models with healthy control as comparison. Tumor tissues were seen in the SAS of both models based on tumor’s morphology, such as darker and denser nucleus with irregular cell and population shape. (A, b) Zoom-in images of different locations (squares), shown in (A, a). (B, a) The fluorescence images of whole brains and spinal cords of the 2 LD models. Green represents MDA-MB231 cells; red represents HDMB03 cells; blue represents DAPI labeled nuclei. (B, b) Zoom-in images of specific regions from (B, a). All the mice were euthanized, followed by profusion of 4% PFA, and decalcification, then sectioned with 12 µm thickness before either H&E staining or fluorescence imaging.

Clinically, LD frequently results in hydrocephalus, which can lead to a range of neurologic sequelae [24]. From a drug delivery perspective, the presence of hydrocephalus is an important consideration that may influence the ability of IT administered therapeutics to access the entirety of tumor metastases across the neuroaxis. A subset of mice bearing LD were therefore subjected to MRI imaging to investigate the size and shape of the cerebral ventricles. MRI images were reconstructed from a transverse perspective, selecting a slice that enabled full visualization of the lateral, 3^rd^, and 4^th^ ventricles (**Figure 5A**). A qualitative increase in size of the ventricles was observed for both BC-LD and MB-LD models (**Figure 5B**). Visualization of the ventricles with MRI confirms evidence from H&E staining that shows an increase in total size of the head, particularly for the BC-LD model that tends to exhibit cranial doming. To estimate ventricular volume quantitively, images were cropped to only include the ventricular system and reconstructed into 3D forms. The size of the ventricular systems was significantly greater than healthy controls for both MB-LD and BC-LD models (**Figure 5C**). MB-LD yielded a 7.3-fold increase and BC-LD a 26.5-fold increase in ventricular volume compared to the healthy control, confirming the presence of substantial hydrocephalus in LD bearing mice.

**Figure 5:**
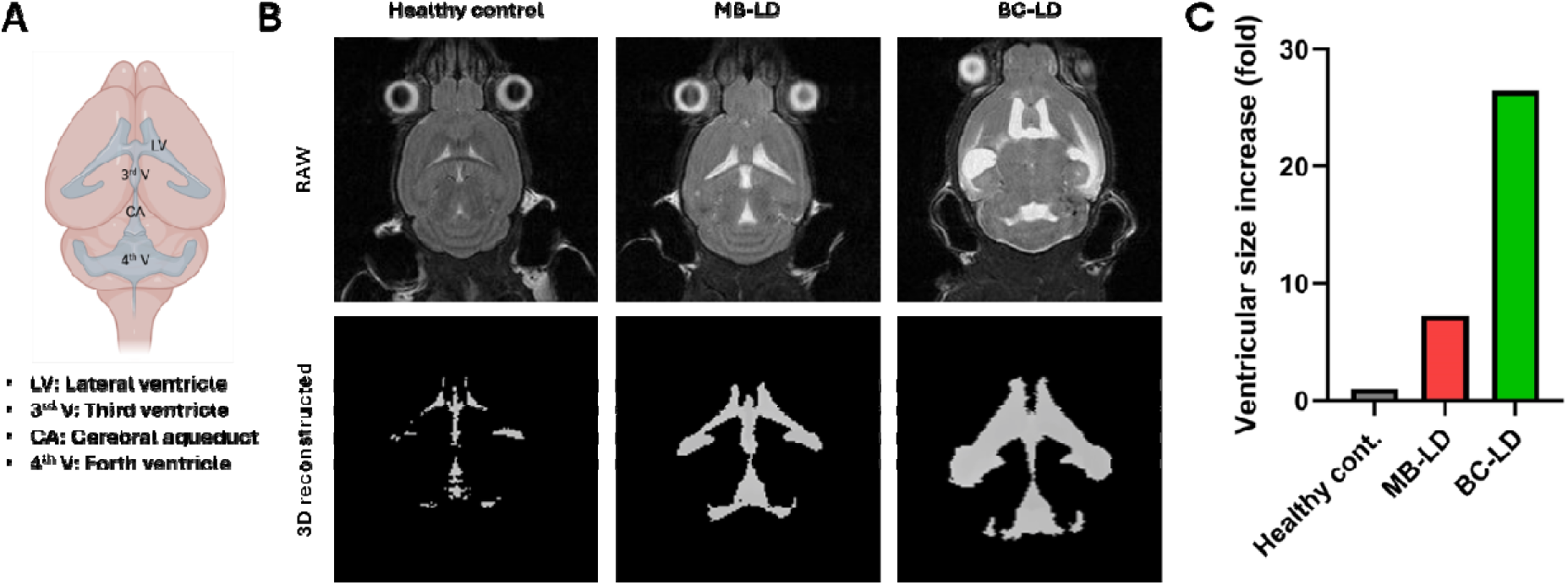
Hydrocephalus was detected in both LD models by 3D imaging using a T2 MRI. (A) Schematic of mouse brain with ventricular systems. (B, top) Raw 2D brain MRI images of healthy, MD-LD model, and BC-LD model. Image location was selected to include lateral ventricle (LV), third ventricle (3^rd^ V), and forth ventricle (4^th^ V). (B, bottom) 3D reconstructed images of the ventricular system. Images were cropped out the only left with ventricles and reconstructed using MATLAB. (C) The size of the ventricular system. The volume was calculated using MATLAB from the 3D reconstructed images at (B, bottom).

### 3.2. Peptide Evaluation

Fluorescently labeled Ang2 and TAT peptides were next administered by the IT-CM route to mice bearing LD, with substance administration occurring once the tumor signal reached 10^9^ photons/sec by IVIS. FITC was used as the fluorescent label for the MB-LD model, while Cy5 was used as the fluorescent label for the BC-LD model; peptides were allowed to circulate for either 2 or 24 hours prior to tissue processing. Here, tumor cell and peptide distribution were assessed across the complete neuroaxis from stereoscopic imaging. As before, tumor was readily detected in the olfactory bulbs, supracerebellar cistern, brainstem, and spinal cord (**Figure 6A**). Peptides were readily detected 2 hours after administration, although the signals dropped substantially by 24 hours. Interestingly, we found that TAT yielded higher signal than Ang2 on for both the MB-LD and BC-LD models, even though these peptides were administered at the same ligand concentration for each model (**Figure 6A**). Quantitative analysis of the of the fluorescence plot profile confirms several important observations: first, that TAT achieves better localization with tumor cells than Ang2 in the MB-LD model, second, that TAT achieves equivalent or better localization with tumor cells than Ang2 in the BC-LD model, and, third, that TAT and Ang2 peptides experienced high levels of clearance or degradation by 24 hours in both models (**Figure 6B**). TAT tended to be detected at a higher, more consistent level in the spinal cord compared to Ang2. While both Ang2 and TAT peptides could be found showing good colocalizing with tumor signal at brain stem and cervical spine, presumably due to the closeness to the injection side, TAT colocalization was improved at distal locations in the spine compared to Ang2. Interesting, after 24 hours of peptide administration, Ang2 and TAT were difficult to detect in both MD- and BC-LD mice. We attribute this to clearance of intact peptide and/or possible degradation of the peptide or peptide label. Although TAT and Ang2 both exhibited reduced signal after 24 hours, TAT signal remained stronger than Ang2 in both LD mice.

**Figure 6.**
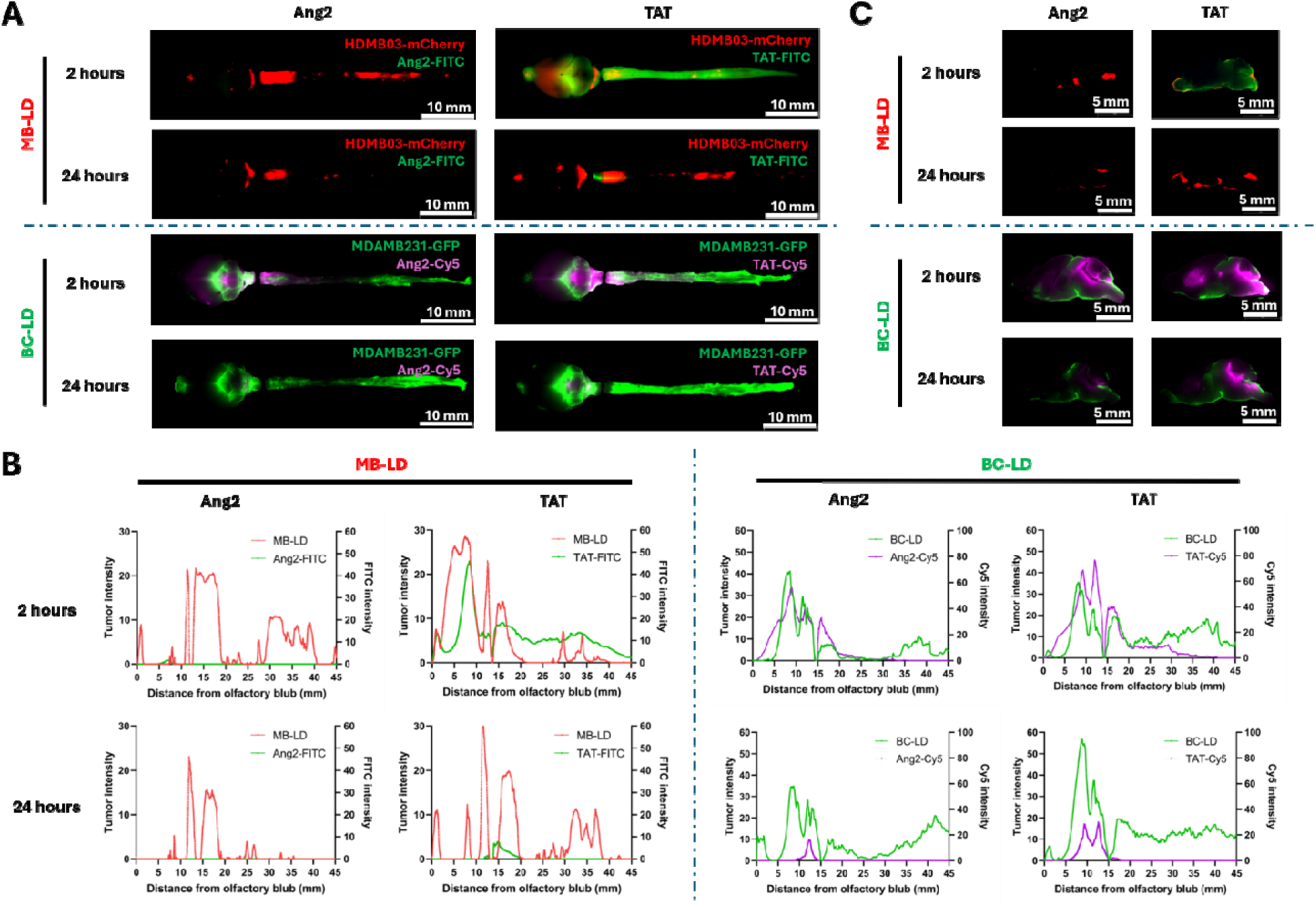
TAT was seen to have better signal than Ang2 throughout out the subarachnoid space in both MB-LD and BC-LD models at 2 hours. (A) Fluorescence images of entire CNS of MB-LD and BC-LD models after 2 and 24 hours of Ang2 and TAT administration via ICM. For MB-LD models, red represents HDMB03-mCherry cells and green represents FITC-conjugated peptides. For BC-LD models, green represents MDA-MB231-GFP cells and purple represents Cy5-conjugatred peptides. All brains and spinal cords were extracted after perfused with 1x DPBS and 4% PFA. (B) Fluorescence images of sagittal section of the brain from (A). All models have tumor signals greater than 10^9^ radiances based on IVIS analysis. All mice subjects were euthanized after 2 or 24 hours of peptide administration. (C) Distribution profiles of tumor and peptide from both MB-LD and BC-LD models. ROI was drawn from olfactory bulbs in (A) in order to compare the distribution of tumor and peptide. The area of the ROI was set the same in order to compare across animal subjects. Blue dash line indicates the separation of brain and spinal cord region.

Following sectioning of the brain into thick slices for fluorescent stereoscopy, we observed that both peptides can reach the entire ventricular system as well as periventricular tissues; TAT in particular was detected in parenchymal regions, including the cerebellum, striatum, and anterior aspects of the brain. Both peptides exhibited strong signal in the cerebellum and brain stem regions, which is the same region in which we detect a high level of LD. However, one important observation from our work is that dissection of the brain and spinal cord out of the skull and vertebral column results in damage and loss of the leptomeninges, including arachnoid mater [25]. Thus, much of the tumor cell population, particularly what has adhered to subarachnoid trabeculae, is expected to be lost upon dissection; this loss is confirmed by our imaging the remaining tumor signal present on the surfaces of discarded bone (**Supplementary Figure 1**). This loss of meningeal signal is particularly important considering that Ang2 is likely to bind to LRP1 that is abundantly expressed in cerebrovascular cells [26]. Indeed, after removing the skin, muscle, and organs, we were able to capture the fluorescence signal of the tumor and peptide through intact bone using a stereoscope (**Supplementary Figure 2**). Similar to the images captured without an axial skeleton, the tumor signals from both MB-LD and BC-LD models were found to be in olfactory bulbs, cerebellum, and throughout the spinal cord into lumbar region (**Supplementary Figure 2**). Thus, extraction of the brain and spinal cord from the skull and vertebral column leads to substantial loss of cell and peptide signal but does not fundamentally shift our understanding of locations of metastasis. It is also important to note that both Ang2 and TAT signals were observed in some locations where tumor signal was not detected, presumably due to the none-specific binding of the peptide to healthy tissue.

To compare the targeting capability of Ang2 and TAT quantitatively, we next performed regional analyses with axial skeleton intact, selecting four regions of interest for deeper: olfactory bulbs, brain stem, thoracic spine, and lumbar spine, where brain stem represents the ventral side of the brain. We observed strong colocalization signals (white arrows) at several spots where TAT peptides potentially bind to the tumor tissues (**Figure 7A**). However, we also witnessed none- significant binding of the peptide to the healthy tissue (for example, non-overlapping signals at thoracic spine and olfactory bulbs, respectively). The non-specific binding to healthy tissue is a known issue for many potential targeting ligands and is most likely due to the non-selective electrostatic interaction of these positively charged peptides interacting with negatively charged cell membranes and/or extracellular matrix. The fact that TAT is more positively charged than Ang2 may contribute to its generally brighter signal across the neuroaxis in both LD models. Interestingly, the brain stem has very strong colocalization signal, presumably because it is close to the administration site, which emphasizes the importance of achieving high exposure of target tissue to targeting ligand. Based on quantitative analysis, targeting achieved by TAT was significantly higher than targeting achieved by Ang2 in thoracic spine for the MB-LD model and in all four tissue regions examined in the BC-LD model (**Figure 7B**). Finally, TAT localization with metastatic cells was examined by confocal microscopy. We found that TAT signal was associated with tumor cells for tissue samples collected from either the brain and spinal cord (**Figure 7C**).

**Figure 7.**
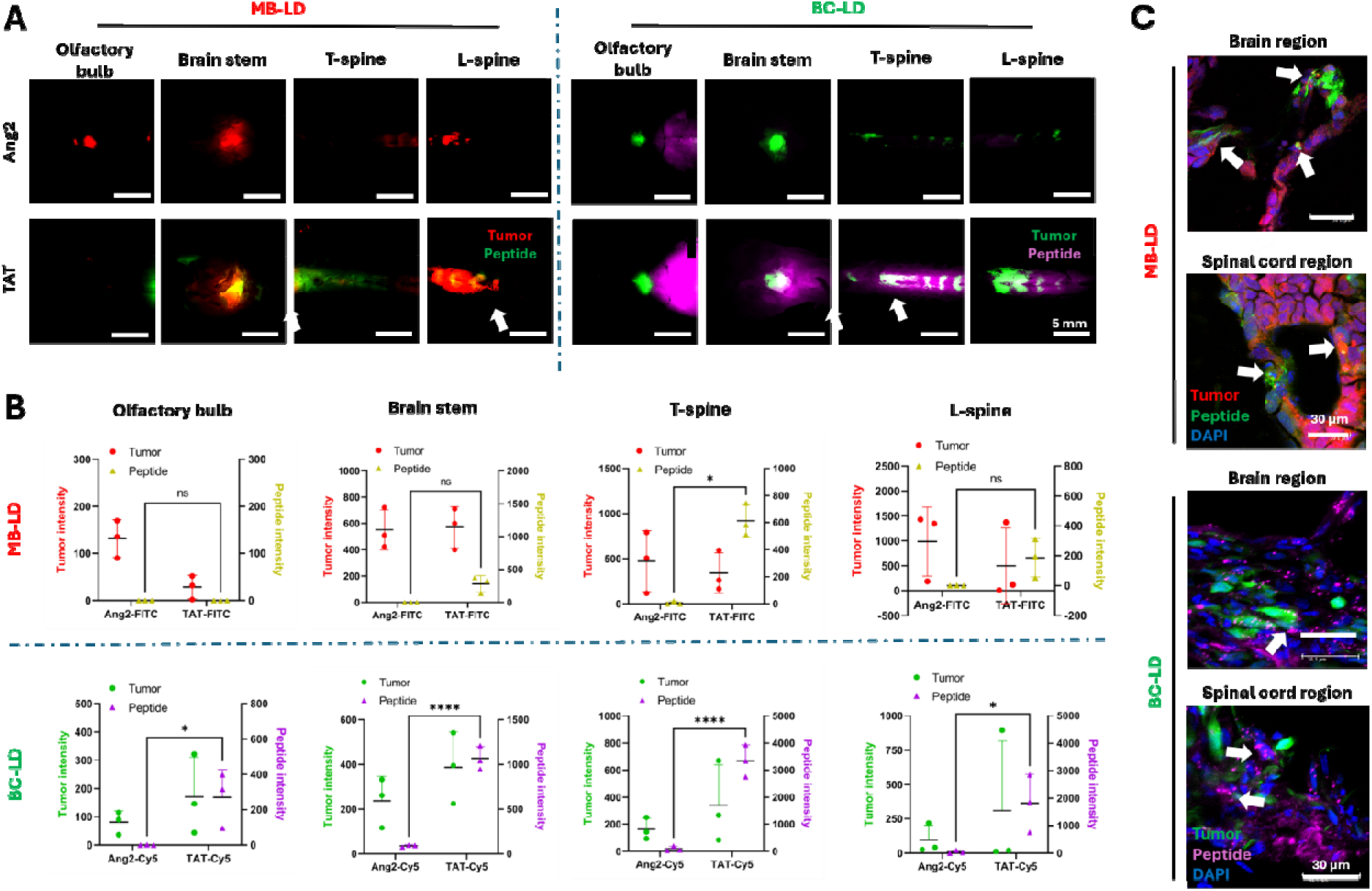
TAT peptide was found at higher level in both MB-LD and BC-LD models at several regions in the CNS and was seen within the tumor cells after tissue sectioning. (A) Regional fluorescence images of the mice after 2 hours of post ICM administration. In MD- LD model, red represents mCherry-labeled HDMB03 cells, green represents peptide signal. In BC-LD model, green represents GFP-labeled MDA-MB231 cells and purple represents peptide signal. White arrow indicates the colocalization of the tumor and peptide signals. (B) Quantitative analysis of TAT binding affinity at desired regions in (a) MD-LD model and (b) BC- LD model. The settings of fluorescence image inquire were different across different regions, but identical in the specific region in order to compare the binding affinity between Ang2 and TAT. (C) Confocal images of the brain and spinal cord tissue sectioned from the mouse administrated with TAT peptide for 2 hours via ICM injection. Tissue was processed with 14 days of decalcification, 2 days of dehydration, OCT embedding, then sectioned at 12 µm thickness. Statistical analysis was carried out using two-way ANOVA with Šidák post hoc test. * Represents p ≤ 0.05; **** represents p ≤ 0.0001 and n=3.

We next examined CNS tropism of TAT and Ang2 after IV administration. Both peptides were poorly detected in the CNS after IV administration; Ang2 results are provided in **Supplementary Figure 3**, and we will focus on TAT results here. TAT was not detected in the CNS either 2 or 24 hours after IV administration, even though the injected peptide mass was around 3 times higher for IV than for ICM injection (**Figure 8A**). By comparing ICM and IV injection routes both at 2- hour time point from both models, ICM route resulted a higher TAT signal than IV route (**Figure 8B**). We further performed the same quantitative analysis to compare the TAT signal at different regions in the CNS via ICM and IV injections. In general, after 2 hours, ICM injection showed the highest signal intensity compared to rest of the conditions ICM after 24 hours, at all regions of the interest. The difference is significantly greater at thoracic spine for MB-LD mice and brain stem, thoracic spine, and Lambor spine for BC-LD mice. However, the signals from both injection methods became almost undetectable at 24-hour time point from both models as well **(Figure 8).** This finding suggests two major observations: first, most of the TAT peptides was potentially washed out or degraded after 24 hrs, even including the peptides that have specifically bound to the tumor tissues (**Figure 6**), and, second, even at longer time point (24- hour), TAT peptide failed to cross the BBB in quantities sufficient for detection. This may highlight the benefits of ICM administration for LD, particularly for initial targeting.

**Figure 8.**
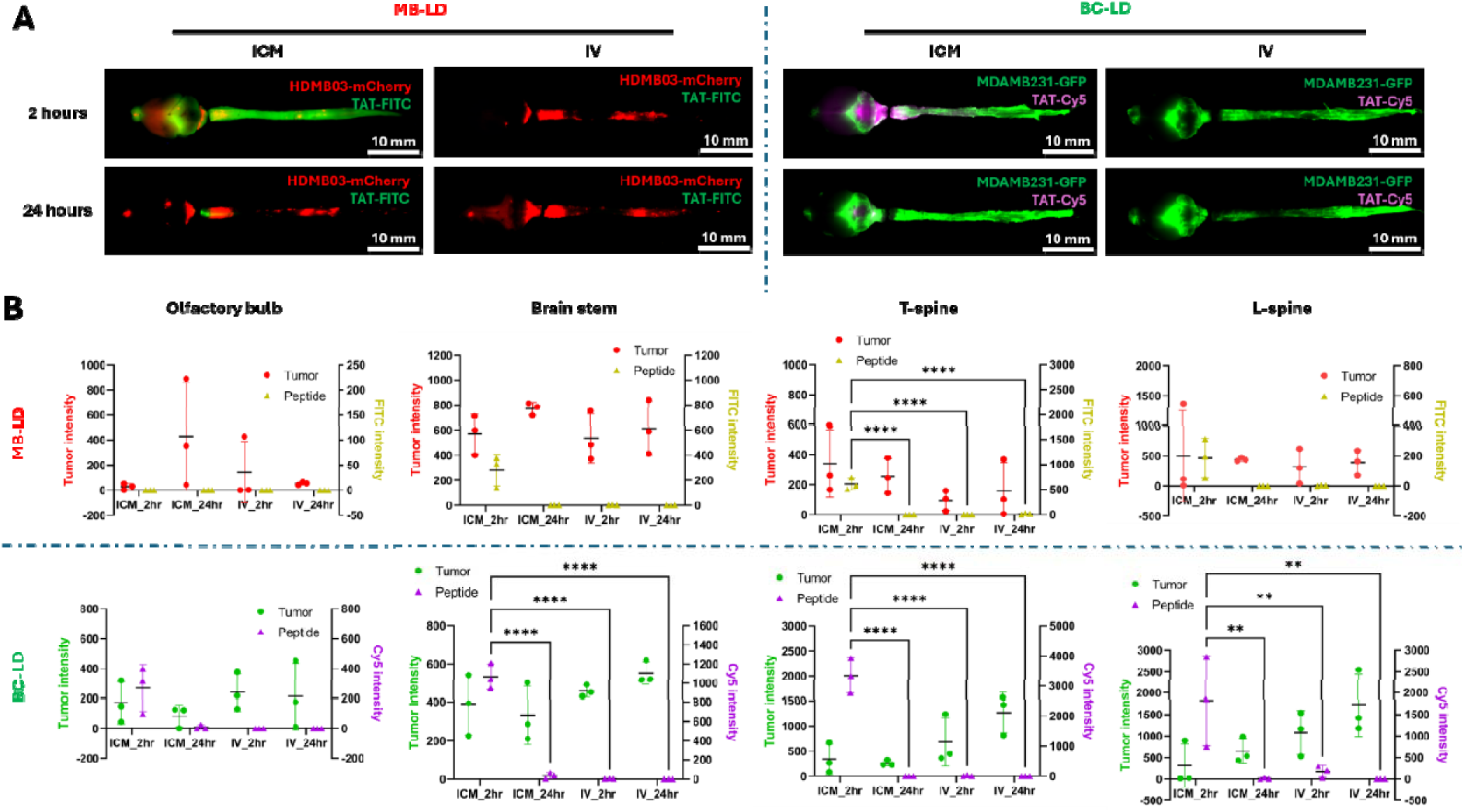
ICM was found to be a better administration route than IV for TAT peptide in both MB-LD and BC-LD models. Qualitative analysis of Ang2 (A) and TAT (B) binding affinity across two different administration routes (ICM vs IV) and two different time points (2 hrs vs 24 hrs) of MB-LD and BC-LD model. (C) Quantitative analysis of TAT binding affinity at four different regions of CNS. The settings of fluorescence image inquire were different across different regions, but identical in the specific region in order to compare the binding affinity across different administration routes and time points. Statistical analysis was carried out using two-way ANOVA with Tukey post hoc test. * Represents p ≤ 0.05, ** represents p ≤ 0.01, **** represents p ≤ 0.0001. n=3 across all conditions.

Peptide uptake was examined *in vitro* with qualitative fluorescence imaging and quantitative flow cytometry analysis. From fluorescence imaging analysis, the results showed that at the same dosage concentration (10 µM), TAT exhibits greater binding to MDA-MB231 cells and HDMB03 cells than Ang2 (**Figure 9A**). The TAT signal could be seen on cellular membrane, cytoplasm, or in the intracellular vesicles within MDA-MB231 cells and HDMB03 cells. To further quantify the difference of the binding, we performed FACS analysis. Both HDMB03 cells and MDA-MB231 cells were incubated with Ang2 and TAT peptides at two concentration (1 and 10 µM) for four time points (0.25, 0.5, 1, and 2 hours). For HDMB03 cells, at 0.25 hours, we observed a slight distribution shift of the cell population dosed with both peptides at both concentrations (**Figure 9B**). This shift was found to be greater when the time point reached 2 hours, suggesting greater binding at longer time point. We also observed that both peptides at 10 µM have greater binding compared to itself at 1 µM, indicating the binding might be concentration dependent. For MDA- MB231 cells, the peak shift was found to be more pronounced than HDMB03 cells while the trend was similar. Two interesting observations follow: first, at all the time points, Ang2 at 10 µM had better binding than TAT at 1 µM; second, multiple peaks were found for the TAT peptide at 10 µM. We attributed this phenomenon to different levels of the peptide association, where some cells might be suddenly saturated with the peptide (right peak) and some cells only took up less (left peak). Therefore, when the time passed, two peaks merged into one. A plot of mean fluorescence intensity (MIF) against time, emphasizes that Ang2 and TAT at both concentrations of 1 µM and 10 µM nearly reached uptake threshold after just 15 minutes of the incubation (**Figure 9B-C**), which was followed by a very slow or no increase until 2 hours of the incubation. This was found in both cell lines and may suggest that 15 minutes or 30 minutes may be efficient for both peptides to associate to the surface of cells and/or enter the cytoplasm. Quantitative analyses (**Figure 9D**) confirm that TAT exhibits a higher degree of binding to both MDA-MB231 and HDMB03 cells compared to Ang2.

**Figure 9.**
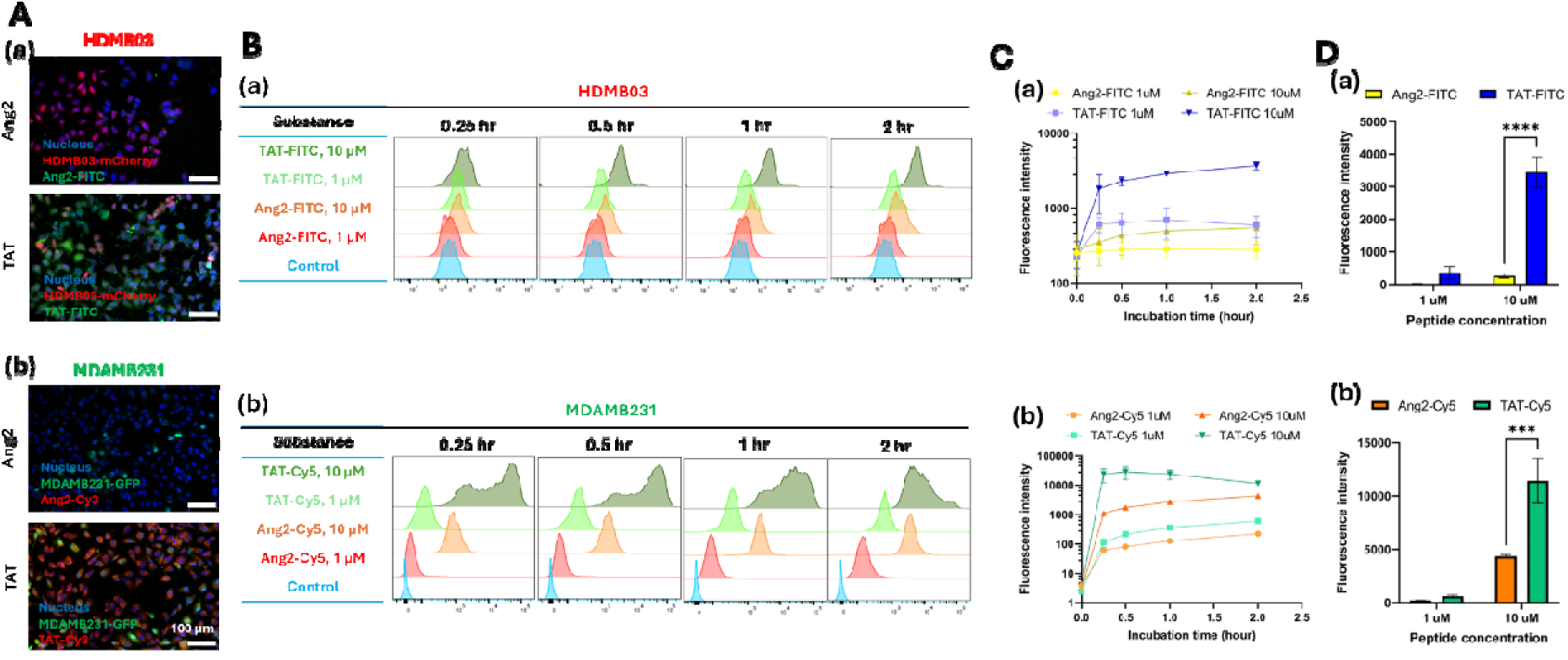
*In vitro* evaluations showed that TAT has significantly higher association towards both MDA-MB231 and HDMB03 cells. (A) *In vitro* cellular uptake images of (a) HDMB03 and (b) MDA-MB231 after 2 hours of peptide incubation. MDA-MB231-GFP cells were incubated with Ang2-Cy3 and TAT-Cy3 while HDMB03-mCherry cells were incubated with Ang2- FITC and TAT-FITC. All cells were incubated with DAPI for nucleus staining at the end. (B) Representative flow cytometry analysis of (a) HDMB03 (b) MDA-MB231. (C) Relative mean fluorescence intensity (MFI) of (a) HDMB03 (b) MDA-MB231 throughout 2 hours incubations of the peptides. (D) The MFI of the peptide at 2 hours, incubated with (a) HDMB03 and (b) MDA- MB231. Statistical analysis was carried out using two-way ANOVA with Tukey post hoc test (n=3). *** Represents p< 0.001; **** represents p< 0.0001.

## 4. Discussion

LD occurs in 5–10% of patients with solid tumors, and it remains a devastating complication of cancer [27]. Several types of tumors that occur in the periphery exhibit a particularly high rate of CNS metastasis, including melanoma, lung cancer, and breast cancer [28]. Approximately 10%- 40% of melanoma patients ultimately developing intracranial involvement [29, 30], and 10%– 36% of the patients with lung cancers are expected to develop LD [31, 32]. Worldwide, approximately 5% of patients with breast cancer will exhibit LD [3, 33], although the rate of LD is higher for TNBC. TNBC is a particularly aggressive form of breast cancer that accounts for 15- 20% of diagnoses, and it is typically less responsive to conventional treatment [34] than other subtypes. Importantly, 25% to 46% of women with TNBC will metastasize to the CNS during the course of their disease [19]. LD can also arise from tumors originating within the CNS, with pediatric MB exhibiting particularly high rates of LD. MB is the most common CNS tumors of childhood, accounting for 10%–15% of pediatric CNS tumors [35]. Group 3 MB accounts for approximately 25% of all cases and carries a particularly poor prognosis, with up to 47% of patients exhibiting LD [20, 36]. LD can be treated with craniospinal irradiation [37], systemic chemotherapy [38, 39], or intrathecally delivered chemotherapy [40–42]; however, the overall survival following a diagnosis of LD is still generally low (<1 year) [27]. Thus, LD remains a highly significant burden on human health.

To address the clinical treatment challenges or to understand the fundamental biology and mechanisms of the disease, experimental brain metastasis models *in vivo* can be extremely valuable. *In vivo* models will allow researchers to recapitulate the architecture and physiology of the given disease in a reproducible setting. Syngeneic models, genetically engineered models, and xenograft models with either primary or immortalized cells have provided significant contributions to the knowledge of brain metastasis pathology and remain pivotal tools for examining novel therapeutic strategies [43, 44]. Among those models, orthotopic xenograft, achieved by transplanting fresh human cancer specimens, patient-derived cells, or immortalized lines, have been shown to better preserve the genomic, histopathological, and phenotypic heterogeneity of the original tissue [43]. However, these models are complicated by the occurrence of a primary tumor (e.g., within the mammary pad or deep in the brain parenchyma) that often drives disease morbidity and mortality with lesser contributions from metastatic disease. Additionally, these models have not been examined from the perspective of drug delivery, i.e., phenotypic considerations for how sites of metastasis are distributed across the neuroaxis. Characterizing these features for distinct LD models was a major goal of our work.

It is already well appreciated that the route of tumor cell inoculation is an important factor in brain metastasis formation [44]. For example, the tumor cells can be placed into tail vein (Intravenous injection), heart (intracardiac injection), carotid artery (intracarotid injection), and specific organ (orthotopic injection). While intracardiac and intracarotid injections can create reliable LD models, tumor dissemination was also seen in other organs [45, 46], a circumstance that fails to capture specific pathophysiology and ligand localization with LD. We therefore applied intrathecal injections to place primary tumor cells directly into SAS [47] to develop models of LD that specifically isolate LD from the primary tumor and without metastasis to other organ sites.

In our study, both LD models developed hydrocephalus by the time mice were moribund (**Figure 5**). We attribute this to the likely obstruction of the CSF outflow from the CNS, where we observed a significant amount of the tumor tissues locating at olfactory bulbs and Interpeduncular cistern (**Figure 4**). These locations are essential for CSF clearance which CSF out flows through ventral side of the brain to the nasal region to reach lymphatic vessels near the nasopharynx before draining to the cervical lymph nodes [48]. The fact that the BC-LD model exhibited a higher degree of hydrocephalus could potentially also be explained by the extensive parenchymal and perivascular infiltration observed in this model, which would be expected to reduce CNS fluid outflow via perivascular routes. Clinically, approximately 1 to 5% patients exhibiting brain metastasis are expected to develop hydrocephalus [49]; the overall survival is only at around 3 months for 50% of the patients even when hydrocephalus is treated aggressively [24, 49]. Tumor-associated hydrocephalus could be the results of the overproduction of CSF, obstruction of the CSF pathways, or reduced CSF absorption which causes ventricular dilatation and increased intracranial pressure [28, 50–52], which may ultimately increase mortality.

Treatment of LD aims to prolong the overall survival, to maintain the quality of patient’s life, and to delay the neurological deterioration. Clinically, IT treatments can be administered through lumbar puncture, cisternal infusion, or ventricular access device (e.g., Ommaya) [53]. Few currently active clinical trials are designed to tackle LD through systemic chemotherapy. For example, a phase II clinical trial was designed to evaluate systemic, intravenous high dose- Methotrexate in breast cancer patients with leptomeningeal metastasis (NCT02422641). Another recent trial was designed to evaluate the efficacy of high dose furmonertinib combined with bevacizumab and pemetrexed (triple therapy with IV administration) for the treatment of non-small cell lung cancer (NSCLC) with leptomeningeal metastasis and epidermal growth factor receptor mutation (EGFRm) through overall survival (OS) (NCT06643000). However, IV administration poses a multitude of challenges, including the presence of the BBB and blood-spinal cord barrier (BCSB), which effectively prevent the vast majority of circulating molecules from reaching the CNS at levels expected to be therapeutically relevant. The BBB and BSCBs are composed of brain microvascular endothelial cells (BMVECs), astrocyte end-feet, and pericytes [54]. This barrier has a significant physiological role as it maintains brain homeostasis and protects the brain from toxins and foreign substances [55], while hindering the therapeutics to reach to the CNS at the same time. In order to cross the BBB, or BSCB, to reach tumors exhibiting LD, various ligand-receptor targeting strategies have been employed, including glucose for the glucose transporter, transferrin for the transferrin receptor, and Ang2 for LDL cholesterol transporters [6]. By conjugating Ang2 with the small molecule chemotherapeutic paclitaxel, ANG1005 has been demonstrated to cross the BBB and was used in several phase I and phase II clinical trials for the treatment of primary brain cancer or cancer that has metastasized to the CNS (NCT00539344, NCT00539383, NCT02048059, NCT01967810). Ang2 has been particularly successful in clinical application; TAT, to our knowledge, has not been used in clinical therapeutics. TAT has been widely used as a targeting ligand in preclinical models. For instance, IV administration of TAT peptide-modified gold nanoparticles led to extensive particle accumulation throughout intracranial metastatic microsatellites [16]. Similarly, Camptothecin-loaded MPEG-PCL-TAT micelles showed significantly prolonged the median survival of intracranial glioma tumor bearing rats [56]. Our data suggest that TAT will be more effective than Ang2 at targeting LD in these two models of cancer metastasis..

In summary, we were able to create two models of metastasis, representing breast cancer and medulloblastoma, that specifically isolate and model LD. Those models carry a unique characteristic which isolates tumor spread within the SAS, enabling us to specifically examine ligand delivery to the SAS and leptomeninges. Our work shows that, with a goal of maximizing ligand localization with target tissues and cells, ICM is preferred over IV, and TAT outperforms Ang2 in these specific tumor models.

---

**Figure S1.**
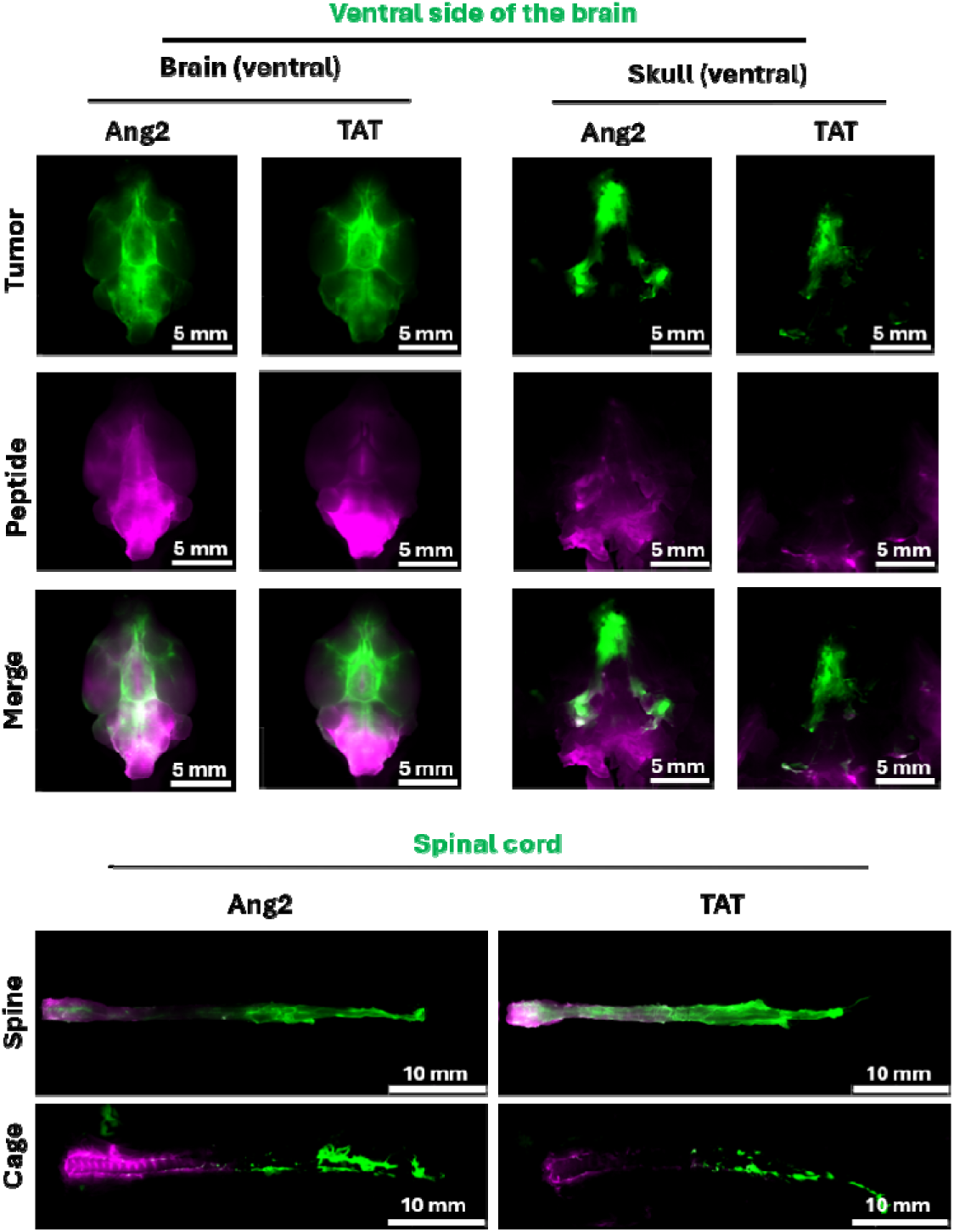
Portions of the tumor and peptide signals were found to remain in the skull and spinal cord cage after the tissue extracted for BC-LD models.

**Figure S2.**
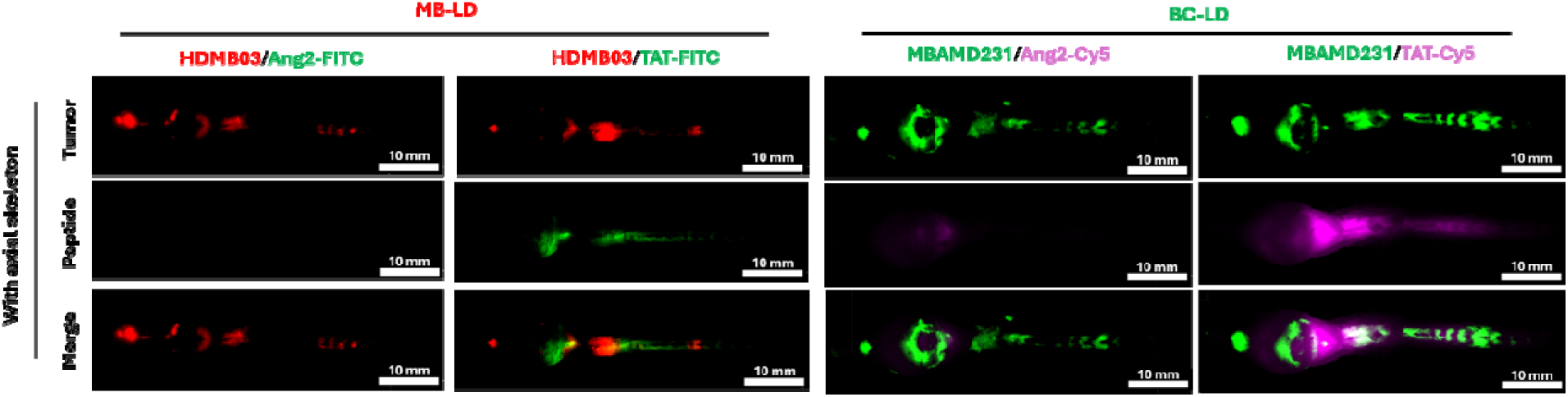
Individual distribution profiles of tumor and peptide from both MB-LD and BC- LD models with axial skeleton.

**Figure S3.**
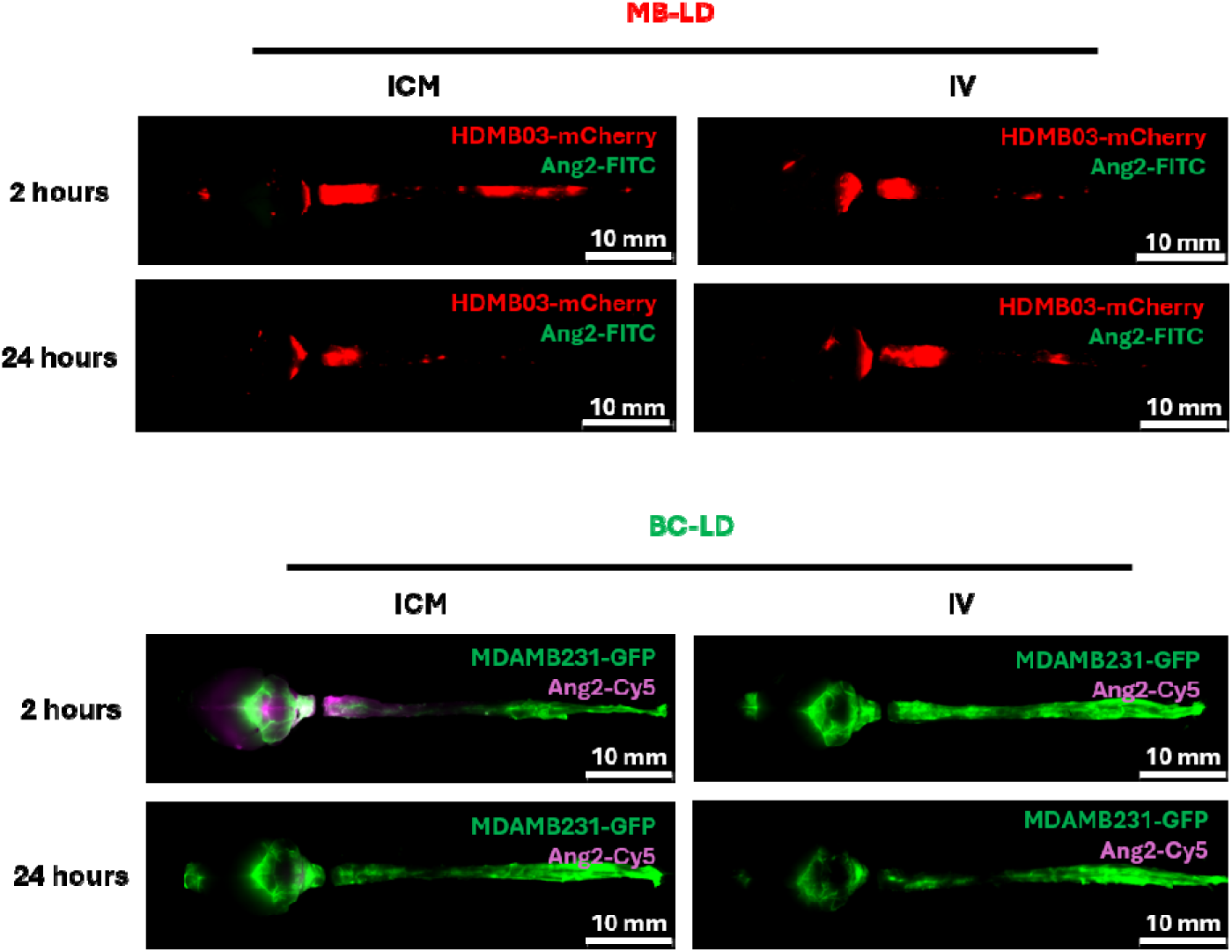
Individual distribution profiles of tumor and Ang2 peptide for both MB-LD and BC-LD models with axial skeleton.

## References

[1] M. Chamberlain, et al., “Leptomeningeal metastasis: a Response Assessment in Neuro-Oncology critical review of endpoints and response criteria of published randomized clinical trials,” Neuro-oncology, vol. 16, no. 9, pp. 1176–1185, 2014.

[2] C. Cocito et al., “Leptomeningeal dissemination in pediatric brain tumors,” Neoplasia, vol. 39, p. 100898, 2023.

[3] S. Hofer and E. Le Rhun, “Leptomeningeal metastases from solid tumours,” memo-Magazine of European Medical Oncology, vol. 14, no. 2, pp. 192–197, 2021.

[4] K. H. Thomas and R. A. Ramirez, “Leptomeningeal disease and the evolving role of molecular targeted therapy and immunotherapy,” Ochsner Journal, vol. 17, no. 4, pp. 362–378, 2017.

[5] D. J. Worm, S. Els-Heindl, and A. G. Beck-Sickinger, “Targeting of peptide-binding receptors on cancer cells with peptide-drug conjugates,” Peptide Science, vol. 112, no. 3, p. e24171, 2020.

[6] Y. Zhao et al., “Recent advances in drug delivery systems for targeting brain tumors,” Drug delivery, vol. 30, no. 1, pp. 1–18, 2023.

[7] Z. Abousalman-Rezvani, A. Refaat, P. Dehghankelishadi, H. Roghani-Mamaqani, L. Esser, and N. H. Voelcker, “Insights into targeted and stimulus-responsive nanocarriers for brain cancer treatment,” Advanced Healthcare Materials, vol. 13, no. 12, p. 2302902, 2024.

[8] P. Kumthekar et al., “ANG1005, a brain-penetrating peptide–drug conjugate, shows activity in patients with breast cancer with leptomeningeal carcinomatosis and recurrent brain metastases,” Clinical Cancer Research, vol. 26, no. 12, pp. 2789–2799, 2020.

[9] A. D. Frankel and C. O. Pabo, “Cellular uptake of the tat protein from human immunodeficiency virus,” Cell, vol. 55, no. 6, pp. 1189–1193, 1988.

[10] I. Ruseska and A. Zimmer, “Internalization mechanisms of cell-penetrating peptides,” Beilstein journal of nanotechnology, vol. 11, no. 1, pp. 101–123, 2020.

[11] R. K. Sahoo, H. Kumar, V. Jain, S. Sinha, Ajazuddin, and U. Gupta, “Angiopep-2 grafted PAMAM dendrimers for the targeted delivery of temozolomide: in vitro and in vivo effects of PEGylation in the management of glioblastoma multiforme,” ACS biomaterials science & engineering, vol. 9, no. 7, pp. 4288–4301, 2023.

[12] X. Cai et al., “Angiopep-2-functionalized lipid cubosomes for blood–brain barrier crossing and glioblastoma treatment,” ACS applied materials & interfaces, vol. 16, no. 10, pp. 12161–12174, 2024.

[13] A. Régina et al., “Antitumour activity of ANG1005, a conjugate between paclitaxel and the new brain delivery vector Angiopep-2,” British journal of pharmacology, vol. 155, no. 2, pp. 185–197, 2008.

[14] Q. Guo et al., “LRP1-upregulated nanoparticles for efficiently conquering the blood-brain barrier and targetedly suppressing multifocal and infiltrative brain metastases,” Journal of controlled release, vol. 303, pp. 117–129, 2019.

[15] B. Fu et al., “Enhanced antitumor effects of the BRBP1 compound peptide BRBP1-TAT-KLA on human brain metastatic breast cancer,” Scientific reports, vol. 5, no. 1, p. 8029, 2015.

[16] R. A. Morshed et al., “Cell-penetrating peptide-modified gold nanoparticles for the delivery of doxorubicin to brain metastatic breast cancer,” Molecular pharmaceutics, vol. 13, no. 6, pp. 1843–1854, 2016.

[17] M. Fowler, J. Cotter, B. Knight, E. Sevick-Muraca, D. Sandberg, and R. Sirianni, “Intrathecal drug delivery in the era of nanomedicine,” Advanced drug delivery reviews, vol. 165, pp. 77–95, 2020.

[18] A. Ruggiero et al., “Intrathecal chemotherapy with antineoplastic agents in children,” Paediatric Drugs, vol. 3, no. 4, pp. 237–246, 2001.

[19] E. Bustamante et al., “Brain Metastasis in Triple-Negative Breast Cancer,” The breast journal, vol. 2024, no. 1, p. 8816102, 2024.

[20] N. Kijima and Y. KaNemura, “Molecular classification of medulloblastoma,” Neurologia medico-chirurgica, vol. 56, no. 11, pp. 687–697, 2016.

[21] C. H. Liu, H. E. D’arceuil, and A. J. De Crespigny, “Direct CSF injection of MnCl2 for dynamic manganese-enhanced MRI,” Magnetic Resonance in Medicine: An Official Journal of the International Society for Magnetic Resonance in Medicine, vol. 51, no. 5, pp. 978–987, 2004.

[22] M.-C. Wu, E. Y. Wang, and T. W. Lai, “TAT peptide at treatment-level concentrations crossed brain endothelial cell monolayer independent of receptor-mediated endocytosis or peptide-inflicted barrier disruption,” PLoS One, vol. 18, no. 10, p. e0292681, 2023.

[23] A. I. Baba and C. Câtoi, “Tumor cell morphology,” in Comparative oncology: The Publishing House of the Romanian Academy, 2007.

[24] N. Lamba, T. Fick, R. Nandoe Tewarie, and M. L. Broekman, “Management of hydrocephalus in patients with leptomeningeal metastases: an ethical approach to decision-making,” Journal of Neuro-oncology, vol. 140, no. 1, pp. 5–13, 2018.

[25] S. A. Grossman and M. J. Krabak, “Leptomeningeal carcinomatosis,” Cancer treatment reviews, vol. 25, no. 2, pp. 103–119, 1999.

[26] T. Kanekiyo, C.-C. Liu, M. Shinohara, J. Li, and G. Bu, “LRP1 in brain vascular smooth muscle cells mediates local clearance of Alzheimer’s amyloid-β,” Journal of Neuroscience, vol. 32, no. 46, pp. 16458–16465, 2012.

[27] J. Remsik and A. Boire, “The path to leptomeningeal metastasis,” Nature Reviews Cancer, vol. 24, no. 7, pp. 448–460, 2024.

[28] N. Wang, M. S. Bertalan, and P. K. Brastianos, “Leptomeningeal metastasis from systemic cancer: review and update on management,” Cancer, vol. 124, no. 1, pp. 21–35, 2018.

[29] E. Tabouret, O. Chinot, P. Metellus, A. Tallet, P. Viens, and A. Goncalves, “Recent trends in epidemiology of brain metastases: an overview,” Anticancer research, vol. 32, no. 11, pp. 4655–4662, 2012.

[30] D. D. Shi et al., “Severe radiation necrosis refractory to surgical resection in patients with melanoma and brain metastases managed with ipilimumab/nivolumab and brain-directed stereotactic radiation therapy,” World Neurosurgery, vol. 139, pp. 226–231, 2020.

[31] J. L. Villano, E. B. Durbin, C. Normandeau, J. P. Thakkar, V. Moirangthem, and F. G. Davis, “Incidence of brain metastasis at initial presentation of lung cancer,” Neuro-oncology, vol. 17, no. 1, pp. 122–128, 2015.

[32] I. T. Gavrilovic and J. B. Posner, “Brain metastases: epidemiology and pathophysiology,” Journal of neuro-oncology, vol. 75, no. 1, pp. 5–14, 2005.

[33] M. Arnold et al., “Current and future burden of breast cancer: Global statistics for 2020 and 2040,” The breast, vol. 66, pp. 15–23, 2022.

[34] N. M. Almansour, “Triple-negative breast cancer: a brief review about epidemiology, risk factors, signaling pathways, treatment and role of artificial intelligence,” Frontiers in molecular biosciences, vol. 9, p. 836417, 2022.

[35] Q. T. Ostrom et al., “Alex’s Lemonade Stand Foundation Infant and Childhood Primary Brain and Central Nervous System Tumors Diagnosed in the United States in 2007–2011,” Neuro-Oncology, vol. 16, no. suppl_10, pp. x1–x36, 2015, doi: 10.1093/neuonc/nou327.

[36] P. A. Northcott et al., “Medulloblastoma Comprises Four Distinct Molecular Variants,” Journal of Clinical Oncology, vol. 29, no. 11, pp. 1408–1414, 2011, doi: 10.1200/jco.2009.27.4324.

[37] J. T. Yang et al., “Randomized phase II trial of proton craniospinal irradiation versus photon involved-field radiotherapy for patients with solid tumor leptomeningeal metastasis,” Journal of Clinical Oncology, vol. 40, no. 33, pp. 3858–3867, 2022.

[38] S. Park et al., “A phase II, multicenter, two cohort study of 160 mg osimertinib in EGFR T790M-positive non-small-cell lung cancer patients with brain metastases or leptomeningeal disease who progressed on prior EGFR TKI therapy,” Annals of Oncology, vol. 31, no. 10, pp. 1397–1404, 2020.

[39] S. M. Tolaney et al., “A phase II study of abemaciclib in patients with brain metastases secondary to hormone receptor–positive breast cancer,” Clinical Cancer Research, vol. 26, no. 20, pp. 5310–5319, 2020.

[40] F. Oberkampf et al., “Phase II study of intrathecal administration of trastuzumab in patients with HER2-positive breast cancer with leptomeningeal metastasis,” Neuro-oncology, vol. 25, no. 2, pp. 365–374, 2023.

[41] I. C. Glitza Oliva et al., “Concurrent intrathecal and intravenous nivolumab in leptomeningeal disease: phase 1 trial interim results,” Nature medicine, vol. 29, no. 4, pp. 898–905, 2023.

[42] E. Le Rhun et al., “Intrathecal liposomal cytarabine plus systemic therapy versus systemic chemotherapy alone for newly diagnosed leptomeningeal metastasis from breast cancer,” Neuro-oncology, vol. 22, no. 4, pp. 524–538, 2020.

[43] J. E. Lee and S. H. Yang, “Advances in brain metastasis models,” Brain Tumor Research and Treatment, vol. 11, no. 1, pp. 16–21, 2023.

[44] I. Daphu et al., “In vivo animal models for studying brain metastasis: value and limitations,” Clinical & experimental metastasis, vol. 30, no. 5, pp. 695–710, 2013.

[45] C. Zhang, F. J. Lowery, and D. Yu, “Intracarotid cancer cell injection to produce mouse models of brain metastasis,” Journal of visualized experiments: JoVE, no. 120, p. 55085, 2017.

[46] J.-K. Pan et al., “ICAM2 initiates trans-blood-CSF barrier migration and stemness properties in leptomeningeal metastasis of triple-negative breast cancer,” Oncogene, vol. 42, no. 39, pp. 2919–2931, 2023.

[47] J. C. Reijneveld, M. J. Taphoorn, and E. E. Voest, “A simple mouse model for leptomeningeal metastases and repeated intrathecal therapy,” Journal of neuro-oncology, vol. 42, no. 2, pp. 137–142, 1999.

[48] Y. Decker et al., “Magnetic resonance imaging of cerebrospinal fluid outflow after low-rate lateral ventricle infusion in mice,” JCI insight, vol. 7, no. 3, p. e150881, 2022.

[49] D. D. Gonda, T. E. Kim, P. C. Warnke, E. M. Kasper, B. S. Carter, and C. C. Chen, “Ventriculoperitoneal shunting versus endoscopic third ventriculostomy in the treatment of patients with hydrocephalus related to metastasis,” Surgical neurology international, vol. 3, p. 97, 2012.

[50] S. H. Lee, D. S. Kong, H. J. Seol, D.-H. Nam, and J.-I. Lee, “Ventriculoperitoneal shunt for hydrocephalus caused by central nervous system metastasis,” Journal of neuro-oncology, vol. 104, no. 2, pp. 545–551, 2011.

[51] E. Pasqualotto, P. H. S. Schmidt, R. O. M. Ferreira, M. P. Chavez, and F. F. S. da Silva, “Endoscopic third ventriculostomy versus ventriculoperitoneal shunt in patients with obstructive hydrocephalus: an updated systematic review and meta-analysis,” Asian Journal of Neurosurgery, vol. 18, no. 03, pp. 468–475, 2023.

[52] D. D. Limbrick Jr and J. R. Leonard, Cerebrospinal fluid disorders: lifelong implications. Springer, 2018.

[53] E. Le Rhun et al., “Diagnosis and treatment patterns for patients with leptomeningeal metastasis from solid tumors across Europe,” Journal of neuro-oncology, vol. 133, no. 2, pp. 419–427, 2017.

[54] P. Ballabh, A. Braun, and M. Nedergaard, “The blood–brain barrier: an overview: structure, regulation, and clinical implications,” Neurobiology of disease, vol. 16, no. 1, pp. 1–13, 2004.

[55] N. J. Abbott, A. A. Patabendige, D. E. Dolman, S. R. Yusof, and D. J. Begley, “Structure and function of the blood–brain barrier,” Neurobiology of disease, vol. 37, no. 1, pp. 13–25, 2010.

[56] H. Taki, T. Kanazawa, F. Akiyama, Y. Takashima, and H. Okada, “Intranasal delivery of camptothecin-loaded tat-modified nanomicells for treatment of intracranial brain tumors,” Pharmaceuticals, vol. 5, no. 10, pp. 1092–1102, 2012.

